# Transferrin receptor is another receptor for SARS-CoV-2 entry

**DOI:** 10.1101/2020.10.23.350348

**Authors:** Xiaopeng Tang, Mengli Yang, Zilei Duan, Zhiyi Liao, Lei Liu, Ruomei Cheng, Mingqian Fang, Gan Wang, Hongqi Liu, Jingwen Xu, Peter M Kamau, Zhiye Zhang, Lian Yang, Xudong Zhao, Xiaozhong Peng, Ren Lai

**Author notes:** These authors contributed equally to this work (Xiaopeng Tang, Mengli Yang, Zilei Duan, Zhiyi Liao, Lei Liu). Corresponding authors (R.L.), (X.P.) or.

## Abstract

Angiotensin-converting enzyme 2 (ACE2) has been suggested as a receptor for severe acute respiratory syndrome coronavirus 2 (SARS-CoV-2) entry to cause coronavirus disease 2019 (COVID-19). However, no ACE2 inhibitors have shown definite beneficiaries for COVID-19 patients, applying the presence of another receptor for SARS-CoV-2 entry. Here we show that ACE2 knockout dose not completely block virus entry, while TfR directly interacts with virus Spike protein to mediate virus entry and SARS-CoV-2 can infect mice with over-expressed humanized transferrin receptor (TfR) and without humanized ACE2. TfR-virus co-localization is found both on the membranes and in the cytoplasma, suggesting SARS-CoV-2 transporting by TfR, the iron-transporting receptor shuttling between cell membranes and cytoplasma. Interfering TfR-Spike interaction blocks virus entry to exert significant anti-viral effects. Anti-TfR antibody (EC50 ~16.6 nM) shows promising anti-viral effects in mouse model. Collectively, this report indicates that TfR is another receptor for SARS-CoV-2 entry and a promising anti-COVID-19 target.

## Introduction

Severe acute respiratory syndrome coronavirus 2 (SARS-CoV-2) which has been assessed and characterized as a pandemic by world health organization on 11 March, 2020 (https://www.who.int/), causes coronavirus disease-19 (COVID-19) with influenza-like manifestations ranging from mild disease to severe pneumonia, fatal acute lung injury, acute respiratory distress syndrome, multi-organ failure, and consequently resulting to high morbidity and mortality, especially in older patients with other co-morbidities^1–10^. As of October 22, 2020, the ongoing COVID-2019 pandemic has swept through 212 countries and infected more than 40 932 220 individuals, posing an enormous burden to public health and an unprecedented effects to civil societies. Unfortunately, to date, there is no vaccine or antiviral treatment for this coronavirus. The pathogenesis and etiology of COVID-19 remain unclear, and there are no targeted therapies for COVID-19 patients ^11^.

Pioneer studies ^12,13^ have demonstrated that angiotensin converting enzyme 2 (ACE2) is the critical receptor for severe acute respiratory syndrome coronavirus (SARS-CoV), which first emerged 17 years ago ^14^. The spike protein of SARS-CoV binds to the host ACE2 receptor and then enters into the target cells. SARS-CoV-2 bears an 82% resemblance to the genomic sequence of SARS-CoV^15^. Especially, the receptor binding domain (RBD) of SARS-CoV-2 is highly similar to the SARS-CoV RBD, suggesting a possible common host cell receptor. Several cryoelectron microscopy (cryo-EM) studies have demonstrated that SARS-CoV-2 spike protein directly binds to ACE2 with high affinity ^16–20^. Soluble ACE2 fused to Ig ^18^ or a nonspecific protease inhibitor (camostat mesylate) showed ability to inhibit infections with a pseudovirus bearing the S protein of SARS-CoV-2 ^21^. Camostat mesylate of high doses (100 mg/mL) has been reported to partially reduce SARS-CoV-2 growth^21^. Recently, Monteil et al., reported that clinical-grade soluble human ACE2 can significantly block early stages of SARS-CoV-2 infections in engineered human tissues ^19^. However, no definite evidence indicates that taking ACEIs/ARBs is beneficial or harmful for COVID-19 infected patients ^11^, suggesting possibilities of other factor/factors assisting in virus entry. Indeed, recent study has shown that neuropilin-1 facilitates SARS-CoV-2 cell entry and infectivity ^22^.

Here, we identified the ubiquitously expressed transferrin receptor (TfR), which is co-localized with ACE2 on cell membranes, as an entry factor of SARS-CoV-2 by directly binding to virus spike protein and ACE2 with high affinities. Moreover, we found that interferences of spike-TfR interaction inhibited SARS-CoV-2 infection in Vero E6 cells and mouse model.

## Results

### Elevated expression of TfR in respiratory tract and lung tissue of monkey and mouse infected by SARS-CoV-2

TfR is ubiquitously expressed ^23^. Given that respiratory tract is a susceptible site for SARS-CoV-2 infection, we employed qRT-PCR and western blot techniques to detect the expression of TfR and ACE2 in several tissues including respiratory tract (nasal cavity, trachea, and lung) and liver. In both RNA and protein levels, the expression of both TfR and ACE2 were significantly elevated in trachea and lung as compared with other tissues (Fig. 1A-C). As illustrated in Fig. 1D and E, TfR was upregulated in lung tissue of SARS-CoV-2 infected monkey and humanized ACE2 (hACE2) mice by immunohistochemical analysis.

**Fig. 1.**
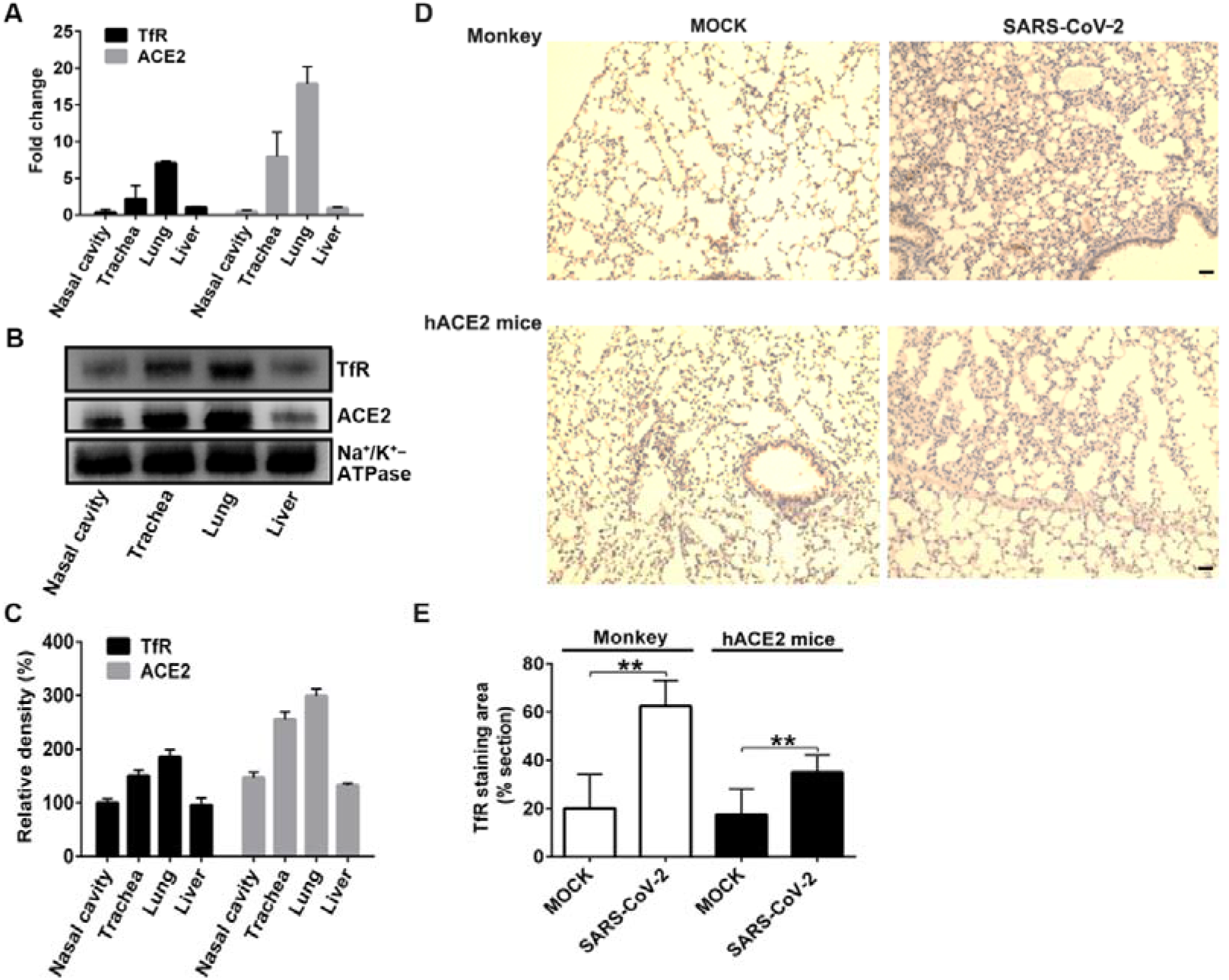
TfR and ACE2 expression are elevated in respiratory tract and lung tissue of monkey and mouse infected by SARS-CoV-2. Transferrin receptor (TfR) and angiotensin-converting enzyme 2 (ACE2) expressions in different tissues including respiratory tract (nasal cavity, trachea, and lung) and liver was detected by qRT-PCR **(A)** and western blot **(B** and **C)**. Na^+^/K^+^ ATPase was used as a control. Immunohistochemical analysis of TfR expression **(D** and **E)** in lung tissue of SARS-CoV-2 infected monkey and d hACE2 mice. Brown cells are positive for TfR. Cell nuclei are stained blue with hematoxylin. Scale bar represents 50 μm.

### Direct interactions among virus spike protein, ACE2 and TfR

Surface plasmon resonance (SPR) was used to study the interaction between TfR and the virus spike protein. As illustrated in Fig. 2A and B, TfR directly interacted with the virus spike protein through enzyme-linked immunosorbent assay (ELISA) and SPR analysis, whereby, the association rate constant (*Ka*), dissociation rate constant (*Kd*), and equilibrium dissociation constant (*KD*) values for the interaction between TfR and spike were 2.69× 10^5^ M^-1^s^-1^, 7.92 × 10^-4^ s^-1^and 2.95 nM, respectively. The effect of TfR on the SARS-CoV-2 spike RBD, which binds to the cell receptor ACE2^24,25^, was also assayed using SPR (Fig. 2C). The *KD* value between TfR and SARS-CoV-2 spike RBD was ~ 43 nM, which is a bit weaker compared with the binding affinity between TfR and SARS-CoV-2 spike.

**Fig. 2.**
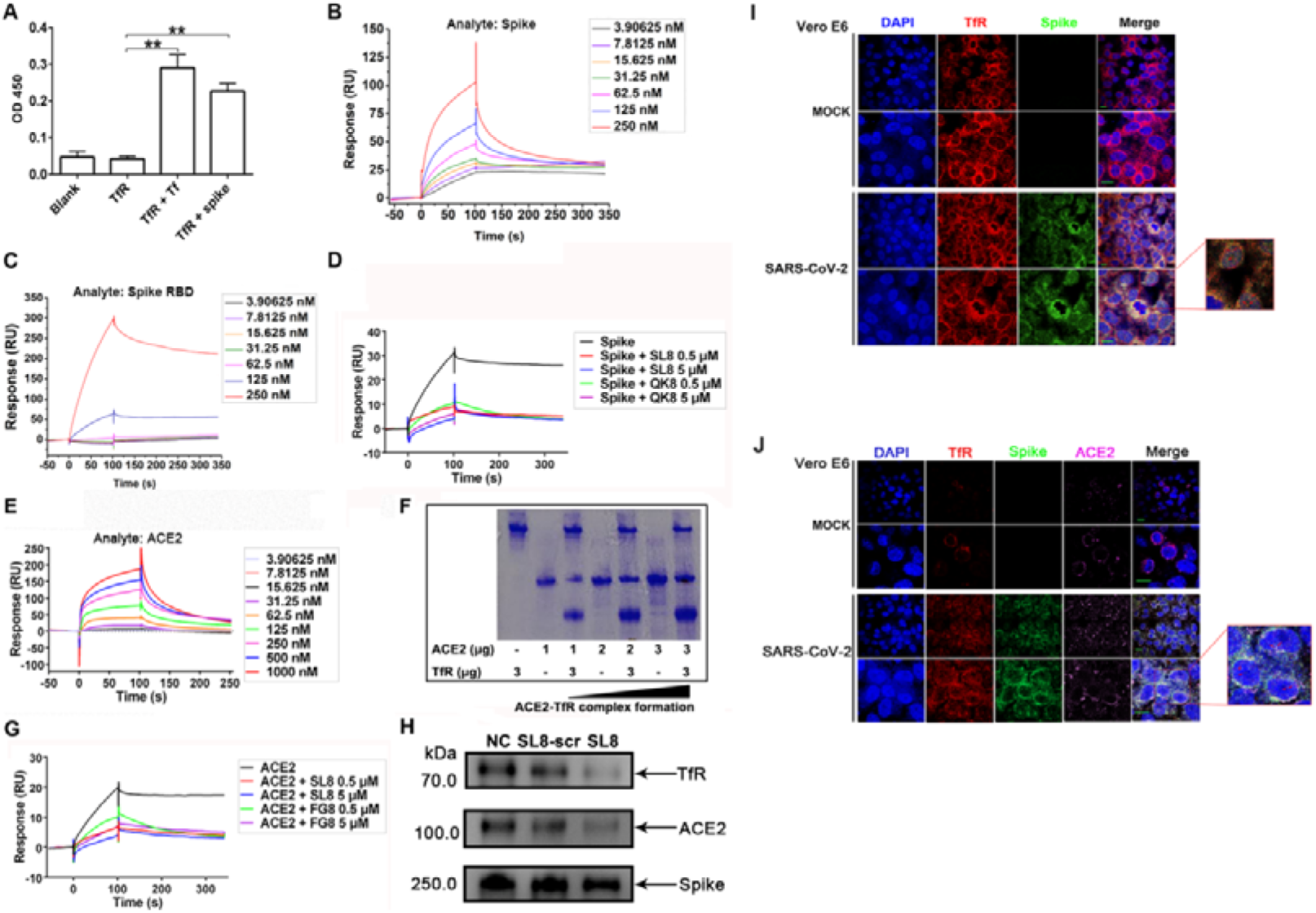
Direct interaction among spike, ACE2 and TfR. **(A)** Interaction between spike and transferrin receptor (TfR) was proved by ELISA. SPR analysis for the interaction between TfR and spike **(B)** or spike receptor binding domain (RBD) **(C)**. **(D)** Effect of inhibitory peptides (SL8 and QK8) on the interaction between spike and TfR by SPR analysis. **(E)** SPR analysis for the interaction between TfR and angiotensin-converting enzyme 2 (ACE2). **(F)** Native gel shift analysis of interaction between TfR (3 μg) and ACE2 (1, 2, and 3 μg). **(G)** Effect of inhibitory peptides (SL8 and FG8) on the interaction between ACE2 and TfR by SPR analysis. **(H)** The effects of SL8 and its scrambled peptide (SL8-scr) on TfR-ACE2-Spike complex formation analyzed by co-immunoprecipitation. Data represent mean ± SD (n = 6), \*\**p* < 0.01 by unpaired t-test **(A)**. Vero E6 cells were infected with SARS-CoV-2 (MOI, 0.2) or uninfected (MOCK) for 2 h. **(I)** Cells were labeled with both anti-transferrin receptor (TfR) and anti-spike antibodies to observe Spike-TfR co-localization. **(J)** Cells were labeled with anti-TfR, anti-angiotensin-converting enzyme 2 (ACE2), and anti-spike antibodies to observe Spike-TfR-ACE2 co-localization. Cell nuclei were labeled by DAPI. White arrows indicate TfR-Spike- or TfR-ACE2-Spike-positive structures on cell membrane. Red arrows indicate TfR-Spike-positive structures in the cytoplasma. Scale bar represents 10 μm. Images are representative of at least three independent experiments.

Based on the TfR structures ^26^ and the virus spike protein ^27^, we made a docking model of TfR-spike interaction (Fig. S1). According to the model, two designed peptides (SL8: SKVEKLTL; QK8: QDSNWASK) were used to interfere with the binding of TfR to spike (Table S1). As illustrated in Fig. 2D, these peptides inhibited TfR-Spike interaction.

As illustrated in Fig. 2E and F, TfR also directly interacted with ACE2 through SPR and native polyacrylamidegel electrophoresis (PAGE) analysis, and the *Ka, Kd*, and *KD* values for the interaction between TfR and ACE2 were 6.33× 10^4^ M^-1^s^-1^, 1.25 × 10^-2^ s^-1^ and 200 nM, respectively. Based on the TfR, ACE2, and spike protein structures, we made docking models of TfR-ACE2 (Fig. S2) and TfR-ACE2-Spike interactions (Fig. S3). According to these models, two inhibitory peptides (SL8 and FG8: FPFLAYSG) were designed to interfere with TfR-ACE2 complex formation (Table S2). As illustrated in Fig. 2G, these peptides inhibited TfR-ACE2 interaction determined by SPR. Notably, co-immunoprecipitation analysis revealed that SL8, but not scrambled peptide of SL8 (SL8-scr), interfered with TfR-ACE2-Spike complex formation (Fig. 2H), indicating that SL8 affected both interactions of TfR-ACE2 and TfR-Spike.

*In vitro* direct interaction between TfR and SARS-CoV-2 has been confirmed as reported above. We next investigated the interaction between TfR and SARS-CoV-2 on cell membranes. As illustrated in Fig. 2I, high density of TfR was found on the membranes of Vero E6 cells. Following the infection of SARS-CoV-2 to the Vero E6 cells, significant co-localization of TfR and SARS-CoV-2 was observed on the cell membranes and in the cytoplasma (Fig. 2I), suggesting that TfR is a membrane receptor for SARS-CoV-2. Further study indicated that TfR was also co-localized with ACE2 on the membranes of both infected and uninfected cells by the virus. Importantly, in the infected cells by the virus, the co-localization of TfR, ACE2, and virus was observed on cell membranes (Fig. 2J), but only TfR-virus co-localization was observed in the cytoplasma, suggesting that the virus is transported into cytoplasma by TfR.

### Interferences of TfR-Spike interaction inhibit SARS-CoV-2 infection

Soluble TfR, Tf, anti-TfR antibody and the designed peptides as mentioned above were used to test their effects on SARS-CoV-2 infections by cytopathic effect (CPE)-based anti-viral assay. As expected, CPE inhibition and quantitative RT-PCR (qRT-PCR) assays indicated that all of them blocked the virus infections to Vero E6 cells (Fig. 3). The concentration to inhibit 50 % viral entry (EC50) determined by CPE assays was 80, 125 and 50 nM (Fig. 3B, D, and F) for soluble TfR, Tf and anti-TfR antibody, while that was 93, 160 and 16.6 nM (Fig. 3C, E, and G) determined by RT-PCR, respectively. There was no cytotoxicity even in their concentration up to 1000 nM (Fig. 3B-G). Notably, the anti-viral effect of 200 nM anti-TfR antibody was comparable to that of high concentration of Remdeivir (4 μM). In addition, the designed peptides showed strong ability to inhibit the viral entry (Fig. 3H-K). At the concentration of 80 μM, SL8, QK8, and FG8 inhibited 87, 99, and 75 % virus infection, respectively (Fig. 3H-K).

**Fig. 3.**
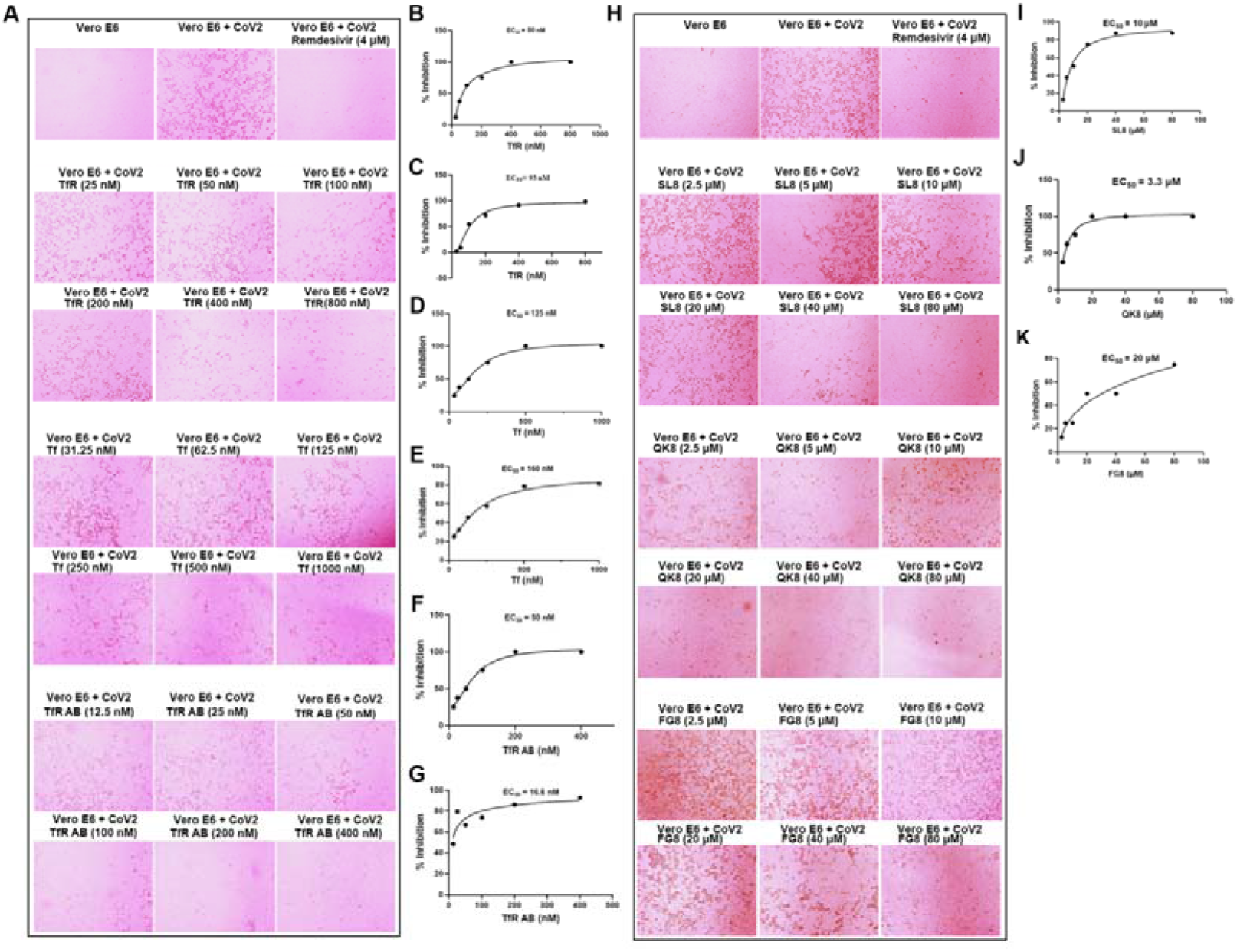
SARS-CoV-2 infection is inhibited through interference of spike-TfR interaction. **(A)** Vero E6 cells were infected with SARS-CoV-2 (MOI, 0.05) in the treatment of different doses of transferrin receptor (TfR, 25, 50, 100, 200, 400, and 800 nM), transferrin (Tf, 31.25, 62.5, 125, 250, 500, and 1000 nM), or transferrin receptor antibody (TfR AB, 12.5, 25, 50, 100, 200, and 400 nM), and representative microscopy images were shown. Their effects on virus entry were evaluated by both quantifying visual CPE read-out **(B, D,** and **F)** and the viral yield in the cell supernatant by qRT-PCR **(C, E,** and **G)**. **(H)** Vero E6 cells were infected with SARS-CoV-2 (MOI, 0.05) in the treatment of different doses of inhibitory peptides (2.5, 5, 10, 20, 40, and 80 μM) for 48 h, and representative microscopy images were shown. Visual CPE **(I-K)** analysis was shown. Remdesivir was used as a positive control. CoV2: SARS-CoV-2.

### SARS-CoV-2 infects ACE2 knockout cells

As illustrated in Fig. 4A and B, ACE2 in Vero E6 and A549 cells were successfully knocked out. Importantly, ACE2-knockout Vero E6 and A549 cells were infected by SARS-CoV-2, and infections in ACE2-knockout Vero E6 and A549 cells were inhibited by anti-TfR antibody (Fig. 4C and D). As illustrated in Fig. S4, the TfR expression levels were first validated by western blot and TfR overexpression promoted virus infection, which was inhibited by TfR down-regulation (Fig. 4E and F).

**Fig. 4.**
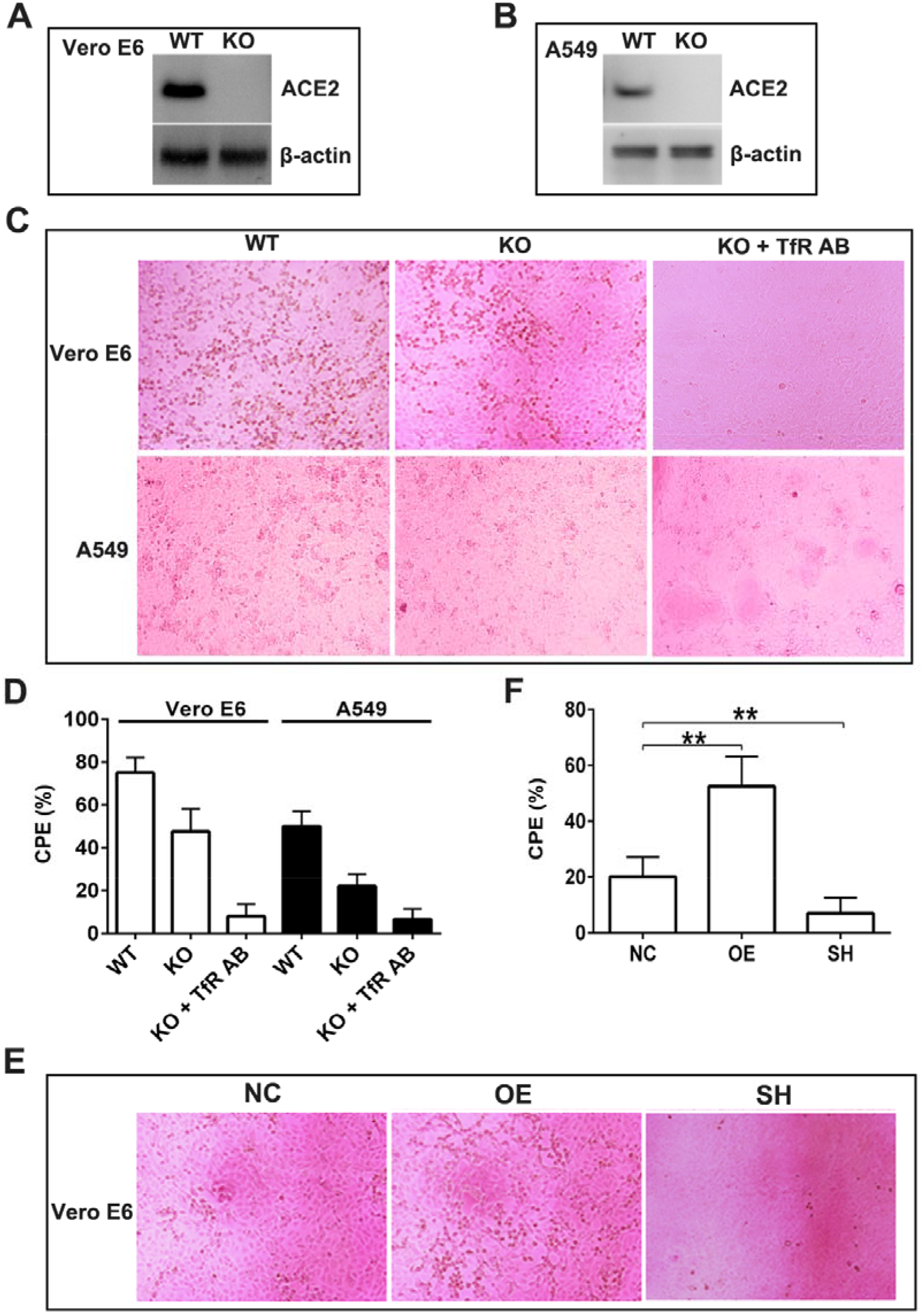
ACE2 knockout cells are infected by SARS-CoV-2. ACE2 expression in wild-type (WT) and ACE2-knockout (KO) Vero E6 **(A)** and A549 **(B)** cells was determined by western blot. **(C)** WT and ACE2-KO Vero E6 cells and A549 cells were infected with SARS-CoV-2 (MOI, 0.2), and representative microscopy images were shown. The effects on virus entry were evaluated by quantifying visual CPE read-out **(D)**. **(E)** TfR over-expression (OE) and knockdown (SH) Vero E6 cells were infected with SARS-CoV-2 (MOI, 0.05), and representative microscopy images were shown. The effects on virus entry were evaluated by quantifying visual CPE read-out **(F)**. Data represent mean ± SD (n = 6), \*\**p* < 0.01 by unpaired t-test **(F)**.

### Humanized TfR (hTfR) mice are sensitized for SARS-CoV-2 infection

The adenoviral vector (AD5) expressing human TfR was constructed as the methods described^28^. Mice were transduced intranasally with Ad5-hTfR, and human TfR expression in lung tissue was validated by western blot (Fig. S5). As illustrated in Fig. 5A, elevated viral replication was detected in lung tissue of hTfR mice at 1, 3, and 5 dpi. Viral infection caused a decrease in mouse body weight, and hTfR showed an obvious decrease in body weight than wild-type mice (Fig. 5B). As illustrated in Fig. 5C, hTfR mice showed more severe vascular congestion and hemorrhage than wild-type mice.

**Fig. 5.**
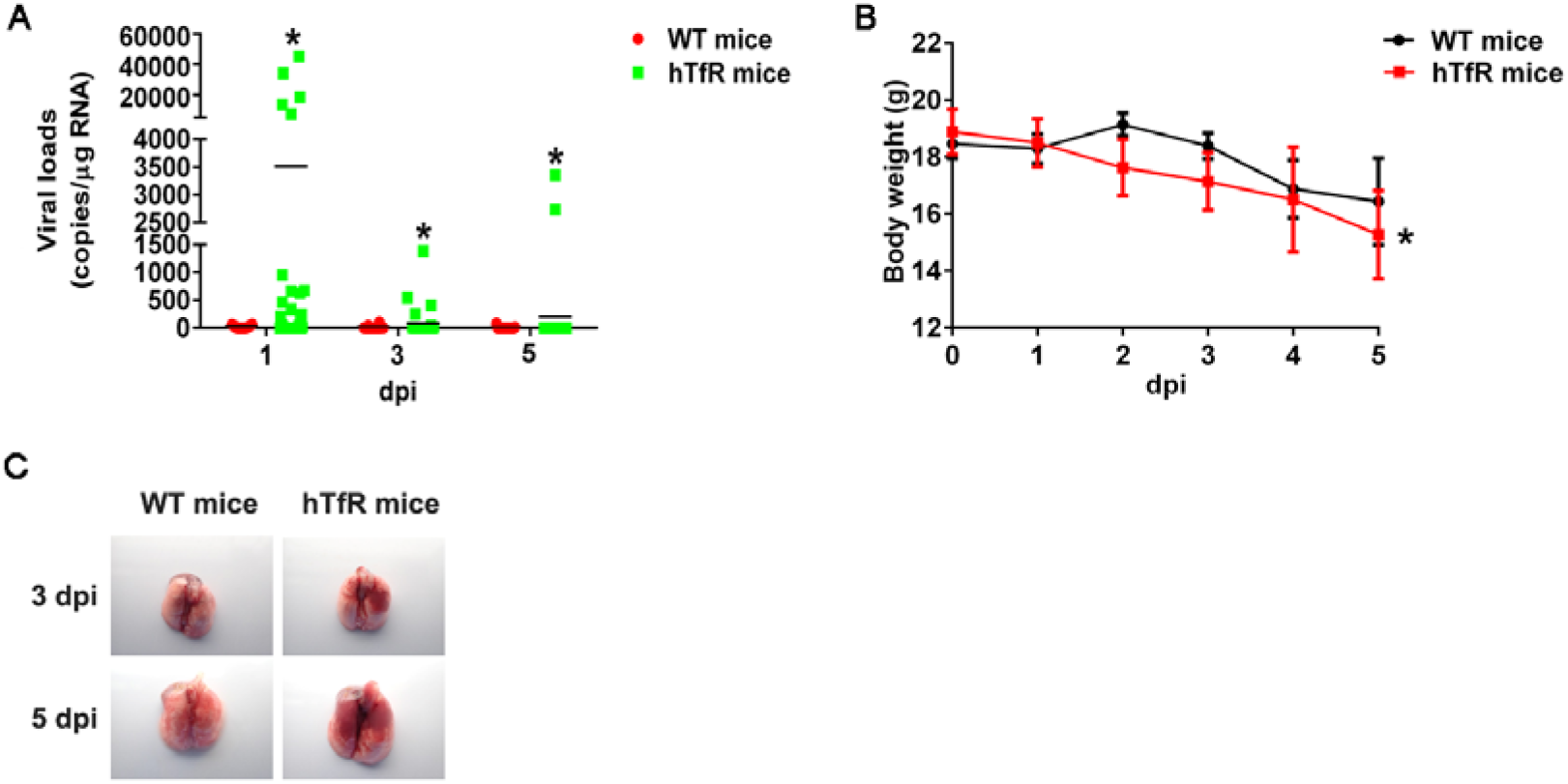
SARS-CoV-2 infects hTfR mice. hTfR mice or wild-type mice (WT mice) were infected with SARS-CoV-2. Lungs of all mice groups were harvested for evaluation of viral loads **(A)** at 1, 3, and 5 dpi, respectively. **(B)** Body weight of all mice groups were recorded. **(C)** Photographs of lung specimens of all mice groups at 3 and 5 dpi are shown. Data represent mean ± SD (n = 6), \**p* < 0.05, \*\**p* < 0.01 by unpaired t-test.

### Anti-TfR antibody shows promising anit-SARS-CoV-2 effects

As illustrated in Fig. 6A, anti-TfR antibody administration inhibited viral replication in mice lung tissue at 3 and 5 dpi, whereas the isotype control IgG administration showed no effects on it. Anti-TfR antibody administration inhibited the decrease in mouse body weight caused by viral infection (Fig. 6B). Histopathological examination of the lungs sections indicated that mice in the control group showed typical interstitial pneumonia (Fig. 6C and D). Anti-TfR antibody administration showed significant protective effects and prevented histopathological injuries caused by virus infection compared with control IgG (Fig. 6C and D).

**Fig. 6.**
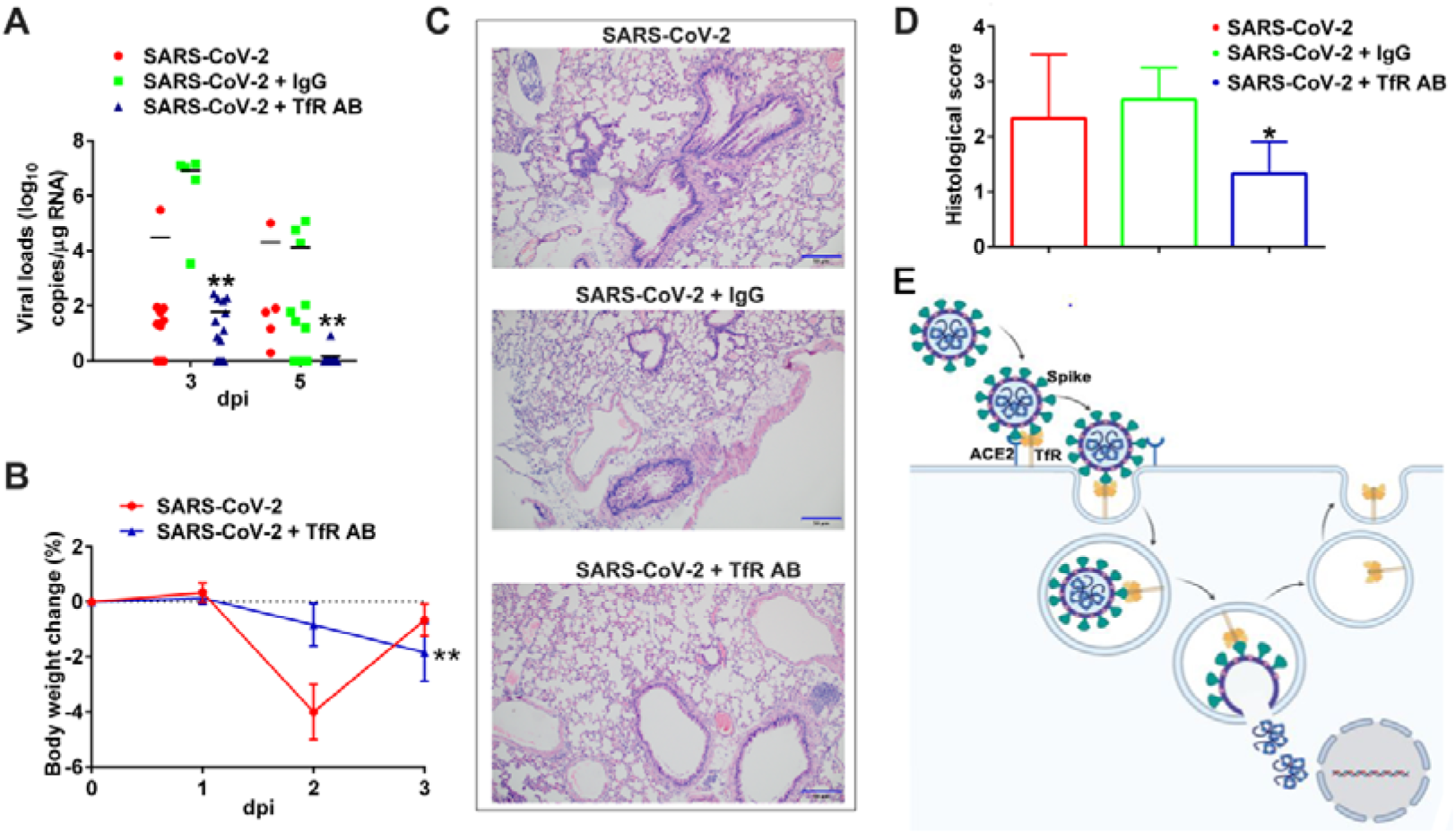
Anti-TfR antibody shows promising anit-SARS-CoV-2 effects. hACE2 mice were infected with SARS-CoV-2. For the TfR antibody (TfR AB) or control IgG-treated group, hACE2 mice were intravenously injected with the TfR antibody or control IgG (1.5 mg/kg) 4 h before SARS-CoV-2 infection. Lungs of all mice groups were harvested for evaluation of viral loads **(A)** at 3 dpi and 5 dpi, respectively. **(B)** Body weight of all mice groups were recorded. **(C** and **D)** Histopathological changes in lungs sections of all mice groups at 5 dpi were analyzed by H & E staining. Scale bar represents 50 μm. Data represent mean ± SD (n = 3), \**p* < 0.01 by unpaired t-test. **(E)** Graphical representation of TfR-ACE2-spike machinery mediating SARS-CoV-2 entry. TfR directly interacts with both ACE2 and virus spike protein to form TfR-ACE2-Spike complex and acts as a machinery of SARS-CoV-2 entry.

## Discussion

The present study reports the identification of TfR, the ubiquitously expressed host receptor on cell membranes, as a new entry factor of SARS-CoV-2 and its elevated expression in respiratory tract upon the virus infetion. TfR mediates SARS-CoV-2 infection by directly binding to spike protein with a high affinity and transporting the virus into host cells. Blocking the interactions among TfR, ACE2 and SARS-CoV-2-Spike protein by using soluble TfR, Tf, anti-TfR antibody, or designed peptides inhibits the virus infection in mouse model, revealing that TfR-Spike complex (also possible TfR-ACE2-Spike complex) is a machinery of SARS-CoV-2 entry and thus providing a new strategies for COVID-19 treatment. Especially, with an EC50 of 16.6 nM, anti-TfR antibody shows promising anti-viral effects in both in vitro and in vivo.

Iron is an essential nutrient element for both host and pathogens. Host innate immune response intensively orchestrates control over iron metabolism to limit its availability during microbe infection. TfR acts as the primary gatekeeper of iron metabolism by binding iron-bound holo-Tf with greater affinity than iron-free apo-Tf and responding to fluctuating iron levels due to the activity of iron response element binding proteins. Cellular iron uptake through circulation to cells is mediated by Tf/TfR system. TfR binds to iron-saturated holo-Tf on the cell surface at pH 7.4 and the TfR-Tf complex is internalized and endocytosed to incorporate Tf-bound ferric ions (Fe^3+^) in endosomes. At a lower pH in the endosomes, Fe^3+^ dissociates from Tf and TfR binds to iron-free apo-Tf without binding to holo-Tf. In the recycling endosomes, the apo-Tf/TfR complex is then transported back to the cell surface where apo-Tf is released into the bloodstream. Both TfR and Tf are reused for another cycle of cellular iron uptake. However, an emerging enigma is that many viruses use the host gate of iron, TfR, as a means to enter into the cells and TfR is a viral target for infection^23,29–31^. The entry and infection of a number different types of viruses including canine parvovirus^32^, mink enteritis virus^33^, feline panleukopenia virus^32, 34, 35^, New World hemorrhagic fever arenaviruses^36,37^, mouse mammary tumor virus^38, 39^, human and simian immunodeficiency viruses^40–42^, hepatitis C virus^43^, human adenoviruses^44^ and alphaviruses^45^ depends on TfR trafficking pathway. Given that TfR is one of the most highly expressed plasma membrane components, different reports suggest that TfR an attractive target for the virus to initiate host cell infection. As illustrated in Fig. 1A-C, an elevated expression level of TfR as well as ACE2 was found in mouse respiratory tract as compared with other tissues. Moreover, TfR is ubiquitously expressed thus increasing the possibility of SARS-CoV-2 infection that is predominantly transmitted among humans via the respiratory route. More importantly, SARS-CoV-2 infected ACE2 knockout cells (Fig. 4A-D), suggesting the presence of another receptor for SARS-CoV-2 entry.

TfR directly binds to spike protein of SARS-CoV-2 with a binding affinity of *KD* 2.95 nM, which falls into a similar range with other binding affinities (31.59-4.67 nM) between ACE2 and SARS-CoV-2 or SARS-CoV as reported by several studies ^16–20^. Compared with the binding affinity between TfR and intact spike protein, a ~15 fold decreased binding between TfR and RBD of SARS-CoV-2 spike protein was found (*KD* of 43 nM), suggesting that other part of the spike protein contributes to the TfR-Spike interaction. The direct TfR-Spike interaction was further confirmed by the colocalization of TfR and the spike protein both on the cell membranes and in the cytoplasma. In addition, TfR directly interacted with ACE2 with a binding affinity of *KD* 200 nM. In both intact and virus-infected cells, TfR was co-localized with ACE2 on cell membranes but the co-localization was not found in the cytoplasma. In virus-infected cells, TfR-ACE2-virus and TfR-virus complex was found on the cell membranes and in the cytoplasma, respectively. No ACE2-virus complex was observed in the cytoplasma. In addition, hTfR mice are sensitized for SARS-CoV-2 infection (Fig. 5). All the data suggest that TfR-Spike interaction mediates SARS-CoV-2 entry on the cell membrane and TfR transports the virus into the cells (Fig. 6E).

As expected, soluble TfR, Tf, anti-TfR antibody or designed peptides significantly blocked SARS-CoV-2’s entry into the Vero cells by interfering with TfR-SARS-CoV-2 interaction thus providing strategies for anti-SARS-CoV-2 treatment and for developing potent therapeutic agents. Especially, the anti-viral effect of 200 nM anti-TfR antibody was comparable to that of high concentration of Remdeivir (4 μM), providing a potential candidate for anti-viral reagent development. According to the docking model (Fig. S1-3), TfR binds to the RBD region of the spike protein interacting with ACE2. The designed peptides (SL8, QK8 and FG8) showed efficacy in inhibiting the virus entry, suggesting an approach to design small molecules of interfering with TfR-SARS-CoV-2 interaction.

## Supplementary material

Supplementary information includes full methods, Fig. S1-5, and Table S1-2.

## Acknowledgments

This work was supported by the National Science Foundation of China (31930015and 21761142002), Chinese Academy of Sciences (XDB31000000, KFJ-STS-SCYD-304, QYZDJ-SSW-SMC012, ZSTH-034, and SAJC201606), and Yunnan Province (2019ZF003, 2018ZF001, and 2019-YT-053) to R.L., as well as the Ministry of Science and Technology of China (2018YFA0801403), National Science Foundation of China (31770835 and 81770464), and Youth Innovation Promotion Association (2017432).

## Author contributions

X.T., M.Y., Z.D., Z.L., L.L., R.C., M.F., G.W., H.L., J.X., P.M., Z.Z., and L.Y. performed the experiments and data analyses; R.L., X.P., and X.Z. conceived and supervised the project; R.L., X.T., and X.P. prepared the manuscript. All authors contributed to the discussions.

## References

1 Munster, V J., Koopmans, M., van Doremalen, N., van Riel, D. & de Wit, E. A Novel Coronavirus Emerging in China - Key Questions for Impact Assessment. New Engl J Med 382, 692–694, doi:10.1056/NEJMp2000929 (2020).

2 Zhou, F. et al. Clinical course and risk factors for mortality of adult inpatients with COVID-19 in Wuhan, China: a retrospective cohort study. Lancet 395, 1054–1062, doi:10.1016/S0140-6736(20)30566-3 (2020).

3 Zhou, P et al. A pneumonia outbreak associated with a new coronavirus of probable bat origin. Nature 579, 270–+, doi:10.1038/s41586-020-2012-7 (2020).

4 Zhu, N. et al. A Novel Coronavirus from Patients with Pneumonia in China, 2019. N Engl J Med 382, 727–733, doi:10.1056/NEJMoa2001017 (2020).

5 Yang, X., Yu, Y & Xu, J. Clinical course and outcomes of critically ill patients with SARS-CoV-2 pneumonia in Wuhan, China: a single-centred, retrospective, observational study (vol 17, pg 534, 2020). Lancet Resp Med 8, E26–E26 (2020).

6 Xu, Z., Shi, L. & Wang, Y Pathological findings of COVID-19 associated with acute respiratory distress syndrome (vol 8, pg 420, 2020). Lancet Resp Med 8, E26–E26 (2020).

7 Lu, R. et al. Genomic characterisation and epidemiology of 2019 novel coronavirus: implications for virus origins and receptor binding. Lancet 395, 565–574, doi:10.1016/S0140-6736(20)30251-8 (2020).

8 Guan, W. J. et al. Clinical Characteristics of Coronavirus Disease 2019 in China. NEngl J Med 382, 1708–1720, doi:10.1056/NEJMoa2002032 (2020).

9 Strabelli, T M. V & Uip, D. E. COVID-19 and the Heart. Arq Bras Cardiol, doi:10.36660/abc.20200209 (2020).

10 Coronaviridae Study Group of the International Committee on Taxonomy of, V. The species Severe acute respiratory syndrome-related coronavirus: classifying 2019-nCoV and naming it SARS-CoV-2. Nat Microbiol 5, 536–544, doi:10.1038/s41564-020-0695-z (2020).

11 Zhang, J., Xie, B. & Hashimoto, K. Current status of potential therapeutic candidates for the COVID-19 crisis. Brain Behav Immun, doi:10.1016/j.bbi.2020.04.046 (2020).

12 Imai, Y. et al. Angiotensin-converting enzyme 2 protects from severe acute lung failure. Nature 436, 112–116, doi:10.1038/nature03712 (2005).

13 Kuba, K. et al. A crucial role of angiotensin converting enzyme 2 (ACE2) in SARS coronavirus-induced lung injury. Nat Med 11, 875–879, doi:10.1038/nm1267 (2005).

14 Drosten, C. et al. Identification of a novel coronavirus in patients with severe acute respiratory syndrome. N Engl J Med 348, 1967–1976, doi:10.1056/NEJMoa030747 (2003).

15 Chan, J. F. et al. Genomic characterization of the 2019 novel human-pathogenic coronavirus isolated from a patient with atypical pneumonia after visiting Wuhan. Emerg Microbes Infect 9, 221–236, doi:10.1080/22221751.2020.1719902 (2020).

16 Walls, A. C. et al. Structure, Function, and Antigenicity of the SARS-CoV-2 Spike Glycoprotein. Cell 181, 281–292 e286, doi:10.1016/j.cell.2020.02.058 (2020).

17 Wan, Y, Shang, J., Graham, R., Baric, R. S. & Li, F. Receptor Recognition by the Novel Coronavirus from Wuhan: an Analysis Based on Decade-Long Structural Studies of SARS Coronavirus. J Virol 94, doi:10.1128/JVI.00127-20 (2020).

18 Wrapp, D. et al. Cryo-EM structure of the 2019-nCoV spike in the prefusion conformation. Science 367, 1260–1263, doi:10.1126/science.abb2507 (2020).

19 Monteil, V. et al. Inhibition of SARS-CoV-2 Infections in Engineered Human Tissues Using Clinical-Grade Soluble Human ACE2. Cell, doi:10.1016/j.cell.2020.04.004 (2020).

20 Lan, J. et al. Structure of the SARS-CoV-2 spike receptor-binding domain bound to the ACE2 receptor. Nature, doi:10.1038/s41586-020-2180-5 (2020).

21 Hoffmann, M. et al. SARS-CoV-2 Cell Entry Depends on ACE2 and TMPRSS2 and Is Blocked by a Clinically Proven Protease Inhibitor. Cell 181, 271–280 e278, doi:10.1016/j.cell.2020.02.052 (2020).

22 Cantuti-Castelvetri, L. et al. Neuropilin-1 facilitates SARS-CoV-2 cell entry and infectivity. Science, doi:10.1126/science.abd2985 (2020).

23 Wessling-Resnick, M. Crossing the Iron Gate: Why and How Transferrin Receptors Mediate Viral Entry. Annu Rev Nutr 38, 431–458, doi:10.1146/annurev-nutr-082117-051749 (2018).

24 Shang, J. et al. Structural basis of receptor recognition by SARS-CoV-2. Nature, doi:10.1038/s41586-020-2179-y (2020).

25 Chen, Y., Guo, Y., Pan, Y & Zhao, Z. J. Structure analysis of the receptor binding of 2019-nCoV. Biochem Biophys Res Commun, doi:10.1016/j.bbrc.2020.02.071 (2020).

26 Lawrence, C. M. et al. Crystal structure of the ectodomain of human transferrin receptor. Science 286, 779–782, doi:10.1126/science.286.5440.779 (1999).

27 Wang, Q. et al. Structural and Functional Basis of SARS-CoV-2 Entry by Using Human ACE2. Cell, doi:10.1016/j.cell.2020.03.045 (2020).

28 Sun, J. et al. Generation of a Broadly Useful Model for COVID-19 Pathogenesis, Vaccination, and Treatment. Cell 182, 734–743 e735, doi:10.1016/j.cell.2020.06.010 (2020).

29 Gkouvatsos, K., Papanikolaou, G. & Pantopoulos, K. Regulation of iron transport and the role of transferrin. Biochim Biophys Acta 1820, 188–202, doi:10.1016/j.bbagen.2011.10.013 (2012).

30 Hentze, M. W., Muckenthaler, M. U. & Andrews, N. C. Balancing acts: molecular control of mammalian iron metabolism. Cell 117, 285–297, doi:10.1016/s0092-8674(04)00343-5 (2004).

31 Sheftel, A. D., Mason, A. B. & Ponka, P. The long history of iron in the Universe and in health and disease. Biochim Biophys Acta 1820, 161–187, doi:10.1016/j.bbagen.2011.08.002 (2012).

32 Parker, J. S. L., Murphy, W. J., Wang, D., O’Brien, S. J. & Parrish, C. R. Canine and feline parvoviruses can use human or feline transferrin receptors to bind, enter, and infect cells. Journal of Virology 75, 3896–3902, doi:Doi 10.1128/Jvi.75.8.3896-3902.2001 (2001).

33 Park, G. S., Best, S. M. & Bloom, M. E. Two mink parvoviruses use different cellular receptors for entry into CRFK cells. Virology 340, 1–9, doi:10.1016/j.virol.2005.06.038 (2005).

34 Palermo, L. M., Hueffer, K. & Parrish, C. R. Residues in the apical domain of the feline and canine transferrin receptors control host-specific binding and cell infection of canine and feline parvoviruses. J Virol 77, 8915–8923, doi:10.1128/jvi.77.16.8915-8923.2003 (2003).

35 Goodman, L. B. et al. Binding site on the transferrin receptor for the parvovirus capsid and effects of altered affinity on cell uptake and infection. J Virol 84, 4969–4978, doi:10.1128/JVI.02623-09 (2010).

36 Radoshitzky, S. R. et al. Transferrin receptor 1 is a cellular receptor for New World haemorrhagic fever arenaviruses. Nature 446, 92–96, doi:10.1038/nature05539 (2007).

37 Sarute, N. & Ross, S. R. New World Arenavirus Biology. Annu Rev Virol 4, 141–158, doi:10.1146/annurev-virology-101416-042001 (2017).

38 Ross, S. R., Schofield, J. J., Farr, C. J. & Bucan, M. Mouse transferrin receptor 1 is the cell entry receptor for mouse mammary tumor virus. P Natl Acad Sci USA 99, 12386–12390, doi:10.1073/pnas.192360099 (2002).

39 Wang, E. et al. Mouse mammary tumor virus uses mouse but not human transferrin receptor 1 to reach a low pH compartment and infect cells. Virology 381, 230–240, doi:10.1016/j.virol.2008.08.013 (2008).

40 Foster, J. L. & Garcia, J.V. HIV-1 Nef: at the crossroads. Retrovirology 5, doi:Artn 8410.1186/1742-4690-5-84 (2008).

41 Koppensteiner, H. et al. Lentiviral Nef suppresses iron uptake in a strain specific manner through inhibition of Transferrin endocytosis. Retrovirology 11, doi:Artn 110.1186/1742-4690-11-1 (2014).

42 Drakesmith, H. et al. HIV-1 Nef down-regulates the hemochromatosis protein HFE, manipulating cellular iron homeostasis. P Natl Acad Sci USA 102, 11017–11022, doi:10.1073/pnas.0504823102 (2005).

43 Martin, D. N. & Uprichard, S. L. Identification of transferrin receptor 1 as a hepatitis C virus entry factor. P Natl Acad Sci USA 110, 10777–10782, doi:10.1073/pnas.1301764110 (2013).

44 Xia, H., Anderson, B., Mao, Q. & Davidson, B. L. Recombinant human adenovirus: targeting to the human transferrin receptor improves gene transfer to brain microcapillary endothelium. J Virol 74, 11359–11366, doi:10.1128/jvi.74.23.11359-11366.2000 (2000).

45 Rose, P. P. et al. Natural Resistance-Associated Macrophage Protein Is a Cellular Receptor for Sindbis Virus in Both Insect and Mammalian Hosts. Cell Host Microbe 10, 97–104, doi:10.1016/j.chom.2011.06.009 (2011).

