## Supplementary Fig. S1-5 and Table S1-2 for "Transferrin receptor is another receptor for SARS-CoV-2 entry"

**Cells and virus**

African green monkey kidney Vero E6 cell line (KCB 92017YJ) was obtained from Conservation Genetics CAS Kunming Cell Bank (Kunming, China) and maintained in in Dulbecco’s modiﬁed Eagle's medium (DMEM, Gibco, USA) containing 10% FBS at 37°C and in the presence of 5% CO_2_.

A clinical isolate SARS-CoV-2 was propagated in Vero E6 cells and amplified SARS-CoV-2 was confirmed via qRT-PCR sequencing and transmission electronic microscopy, and titrated via plaque assay (10^6^ pfu/ml). Infection experiments were performed in a biosafety level-3 (BLS-3) laboratory of National Kunming High-level Biosafety Primate Research Center, Yunnan China.

**Animals and ethics statement**

All animal experiments were approved by the Animal Care and Use Committee of the Kunming Institute of Zoology (SMKX-20200627-08) and conformed to the US National Institutes of Health’s Guide for the Care and Use of Laboratory Animals (National Academies Press, 8th Edition, 2011). Specific-pathogen-free (SPF) C57BL/6J mice (male, 8 weeks old,10 back crosses) were purchased from the Institute of Laboratory Animal Sciences, Chinese Academy of Medical Sciences. All mice were housed under a 12 h light-12 h dark cycle at 24°C and tested at 10 weeks of age.

**Tissue distribution of transferrin receptor (TfR)**

Organs including respiratory tract (nasal cavity, trachea, and lung) and liver were collected from C57 mice and placed in RNAlater for RNA extraction or PBS for protein extraction. The RNA extraction and cDNA reverse transcription procedures were performed by using an RNA extraction kit (DP419, Tiangen, China) and reverse transcription kit (A5000, Promega, USA), respectively, as per the manufacturer’s instructions. TfR and angiotensin-converting enzyme 2 (ACE2) expression was quantified by both quantitative RT-PCR (qRT-PCR) (forward primer (5’-3’): GAGGAACCAGACCGTTTAGTTGT and reverse primer (5’-3’): CTTCGCCGCAACACCAGCA for TfR; forward primer (5’-3’): ATATGACTCAAGGATTCTGGG and reverse primer (5’-3’): GCTGCAGAAAGTGACATGATT for ACE2) and western blot. PCR was performed on a CFX-96 Touch Real-Time Detection System (Bio-Rad, USA). Western blot was performed as previous described^1^. Briefly, tissue homogenates were first separated by a 12% sodium dodecyl sulfate-polyacrylamide gel electrophoresis (SDS-PAGE) and then transferred to polyvinylidene difluoride (PVDF) membranes. Anti-TfR (1:2000, ab1086, Abcam, USA) and anti-ACE2 (1:2000, BS90021, Bioworld, China) antibodies were used for immunoreactivity. To control the loading and transfer, blotting for Na+/K+ ATPase was used as a loading control. TfR in lung tissue of SARS-CoV-2 infected monkey and humanized ACE2 (hACE2) mice were also stained using standard immunohistochemistry techniques as per the methods described^2^.

**Enzyme-linked immunosorbent assay (ELISA)**

In brief, TfR (10 μg/ml; 11020-H07H, Sino Biological, China) was diluted with coating buffer (0.01 M sodium carbonate, pH 9.6) and adsorbed to 96 well flat-bottom immunoplates (Nunc, Roskilde, Denmark; 100 μl per well) at 4°C overnight. Plates were then blocked with 1% bovine serum albumin (BSA, 36101ES7, YEASEN, China) in a phosphate-buffered saline (PBS) with 1% Tween-20 (v/v)) for 1 h at 37 °C. After washing the immunoplates with the wash solution for 3 times, spike (10 μg/ml) was added and incubated for 1 h at 37°C. The first anti-spike antibody (1:5000 dilution, 40150-R007, Sino biological, China) in the wash solution was incubated for 1 h at 37°C after washing the immunoplates with the wash solution for 3 times. After washing the immunoplates with the wash solution for 3 times, a second anti-rabbit IgG antibody (1:5000 dilution, horseradish peroxidase (HRP) labeled, KPL, USA) was then added and incubated for another 1 h at 37°C. Color development was carried out with 100 μl 3, 3’, 5, 5’-tetramethylbenzidine (TMB, PR1200-500ml, Solarbio, China), and the reaction was stopped by stop solution (1 M H_2_SO_4_). The absorbance at 450 nm was monitored on a plate reader.

**Surface plasmon resonance (SPR) analysis**

BIAcore 2000 (GE, USA) was used to analyze the interaction among TfR, spike, spike receptor binding domain (RBD) and angiotensin-converting enzyme 2 (ACE2). Briefly, TfR was first diluted (20 μg/ml) with 200 μl of sodium acetate buffer (10 mM, pH 5) and then flowed across the activated surface of a CM5 sensor chip (BR100012, GE, USA) at a flow rate of 5 μl/min, reaching a resonance unit (RU) of ~2000. The remaining activated sites on the chip were blocked with 75 μl of ethanolamine (1 M, pH 8.5). Serial concentrations of spike protein (3.90625, 7.8125, 15.625, 31.25, 62.5, 125, 250 nM; Z03481, Genscript, USA), spike RBD (3.90625, 7.8125, 15.625, 31.25, 62.5, 125, 250 nM; Z03479, Genscript, USA), or ACE2 (3.90625, 7.8125, 15.625, 31.25, 62.5, 125, 250, 500, and 1000 nM; 10108-H08H, Sino biological, China) in Tris-HCl buffer (20 mM, pH 7.4) were applied to analyze their interactions with immobilized TfR at a flow rate of 10 μl/min. The *KD* for binding as well as the *Ka* and *Kd* rate constants were determined using the BIA evaluation program (GE, USA). Spike (60 nM) or ACE2 (60 nM) mixed with different concentrations of peptides (0.5 and 5 μM) was applied to analyze the blockage of designed peptides at the interaction between spike or ACE2 and TfR at a flow rate of 20 μl/min.

**Native PAGE**

Blue Native Polyacrylamide Gel Electrophoresis (BN PAGE) technique was used to analyze interaction between TfR or ACE2 as per the manufacturer’s protocols (NativePAG™Novex Bis-Tris Gel System, BN1001BOX, Life technologies, USA).

**Immunoprecipitation**

Spike (2 μg), TfR (2 μg), and ACE2 (2 μg) were mixed with or without designed peptides (1 μM) for 30 min at 4 °C, and then anti-spike antibody (5 μg, 40150-R007, Sino biological, China) was added and incubated for 16 h at 4 °C in 30 μl of Tris-HCl buffer (25 mM, pH 7.4). Protein A agarose (20 μl, P2006, Beyotime, China) was then added and incubated for 3 h at 4 °C. After centrifugation at 2 500 rpm for 5 min at 4 °C, loading buffer (10 μl, 4 × CW0027A, CWBIO, China) was added, followed by boiling for 10 min to obtain the coupled proteins. All proteins were subjected to 12% SDS-PAGE separation and polyclonal antibodies against spike, TfR, and ACE2 (1:2000 dilution, BS90021, Bioworld, China) were used to identify spike, TfR, and ACE2, respectively.

**Protein-protein docking**
To model the SARS-CoV-2-spike-TfR complex, we used the known crystal structure of TfR (PDB ID: 1CX8) and SARS-CoV-2-spike (PDB ID: 6LZG) protein for protein docking. After running a short time (10 ps) Molecular Dynamics (MD) simulation to obtain the optimized structure model, the SARS-CoV-2-spike was docked to TfR by ZDOCK. Protein-protein docking was guided by SPR experimental data (Fig. 2B and C), where residues that disrupted binding were forced to be included in the interface, and residues that did not affect binding were forced not to be included. About 2000 structure complexes were generated and ranked according to the ZRANK scoring function. The best ZDOCK pose between these two conformations was therefore used as a representative of SARS-CoV-2-spike-TfR interactions and verified (Fig. S1). For modeling the TfR-ACE2 complex, the same PDB files were used for docking. Docking results were guided by SPR, (Fig. 2E). About 10 ns MD simulation was used for optimizing the complex model (Fig. S2).

**Confocal microscopy**

To colocalize TfR, spike protein, and/or ACE2 at the membrane surface and the cytoplasma of Vero E6 cells, cells were infected with SARS-CoV-2 (MOI, 0.2) or uninfected (MOCK) for 2 h. After washing with PBS, cells were fixed for 15 min with 4% paraformaldehyde in PBS, blocked for 1 h at room temperature with 1% BSA, and then incubated with the antibodies against TfR (1:200 dilution, 11020-MM04, Sino Biological, China), spike protein (1:200 dilution, 40150-R007, Sino biological, China), and ACE2 (1:400, AF5165, Affinity, USA) for 1 h at 37 °C. After washing for three times with PBS to remove the excess primary antibody, the section was incubated with a fluorescently labeled secondary antibodies for 1 h at 37 °C. Following washing with PBS to remove excess secondary antibodies, cells were stained with Prolong Gold Antifade DAPI (P36941, Life Technologies, USA) and imaged with a confocal microscope (FluoView™ 1000, Olympus, USA).

**Antiviral activities evaluation**

To evaluate the antiviral efficacy of TfR, transferrin (Tf; T4382, Sigma, USA), TfR antibody (ab1086, Abcam, USA), or interference peptides, Vero E6 cells were pre-treated with the different doses of these samples for 1 h, and virus (MOI, 0.05) was subsequently added to allow infection for 1 h. Then, the virus-protein mixture was removed and cells were further cultured with fresh protein-containing medium. At 48 h, further cytopathogenic effect (CPE) was analyzed and cell supernatant was collected and lysed in lysis buffer (15596018, Thermo, USA) for qRT-PCR quantification analysis as the methods described previously^3,4^. For qRT-PCR, the protocol was performed as described above. Primers specific for NP gene was, Target-2-F: GGGGAACTTCTCCTGCTAGAAT, Target-2-R: CAGACATTTTGCTCTCAAGCTG, Target-2-P: 5'-FAMTTGCTGCTGCTTGACA GATT-TAMRA-3' as described previously^5^.

**CRISPR/Cas9-mediated ACE2 knockout**

We used the CRISPR/Cas9 system to establish stable ACE2 knockout in Vero E6 and human A549 cell. For this study, we designed sequences targeting ACE2 locus of Vero E6 (AGCCTGGCTCGGCAAGTAGA) and human A549 cell (CAAGTGAACTTTGATAGAAC), respectively. The guide RNA (gRNA) sequences for ACE2 were designed and cloned into the lentiCRISPRv2 puro plasmid. Constructed plasmids were co-transfected into HEK293T cells with the packaging plasmids pCMVD8.9 and pMD2.G at a ratio of 10:5:2. The culture medium was collected and filtered. Then using the supernatant containing the lentiviral particle infected Vero E6 or A549 cells. After 24 hours of puromycin screening, single-cell culture was established using the limiting dilution method. The ACE2-null clones were screened by DNA and cDNA sequencing and western blotting.

**TfR overexpression and knockdown**

The coding region of the green monkey TfR (GenBank: 103242011) was synthesized and cloned into pLVX-Puro lentiviral plasmid (Clontech, USA) as per the method of our previous manuscript^1^. The oligonucleotides of the shRNA sequence targeting green monkey TfR sequence (CGTGAATTTAAACTCAGCAAA) was synthesized by Sangon Biotech (Shanghai, China) and was inserted into the BamHI and EcoRI sites of the RNAi-Ready pSIREN-RetroQ retroviral vector (Clontech, USA) as as per the method of our previous manuscript^1^. The lentiviral vector for TfR overexpression and retroviral vector for TfR knockdown were transfected into Vero E6 cells to construct TfR overexpression or knockdown cells.

**Human TfR expression adenoviral vector construction and transduction of mice**

The studies were performed in an animal biosafety level 3 (ABSL3) facility, and approved by the Institutional Committee for Animal Care and Biosafety in the Kunming Institute of Zoology (China). All experiments complied with all relevant ethical regulations. The adenoviral vector AD5 expressing hTfR was constructed as the methods described^6^. Mice were anesthetized and transduced intranasally with 3 × 10^8^ FFU of empty Ad5 or Ad5-hTfR in DMEM. Mice were infected intranasally with 2 × 10^6^ TCID50 SARS-CoV-2 after seven days transduction. The infected mice were observed daily to record body weight and temperature. The mice were euthanized and dissected at 1, 3, and 5 days post-infection (dpi), respectively, to collect different tissues to screen virus replication and histopathological changes.

**Anti-SARS-CoV-2 activities in hACE2 mice**

Male hACE2 mice were provided by the Guangzhou Institutes of Biomedicine and Health (China). After anesthetization, hACE2 mice were inoculated intranasally with SARS-CoV-2 stock virus at a dose of 2 × 10^6^ TCID50. For the TfR antibody or control IgG-treated groups, hACE2 mice were intravenously injected with the TfR antibody or control IgG (1.5 mg/kg) 4 h before SARS-CoV-2 infection. The infected mice were observed daily to record body weight and temperature. The mice were euthanized and dissected at 1, 3, and 5 days post-infection (dpi), respectively, to collect different tissues to screen virus replication and histopathological changes.

**Statistical analysis**

The data obtained from independent experiments are presented as means ± SD. All statistical analyses were two-tailed with 95% confidence intervals (CI). The results were analyzed using unpaired t-test by Prism 6 (GraphPad Software) and SPSS (SPSS Inc, USA). Differences were considered significant at p < 0.05.

**Supplementary Figures**

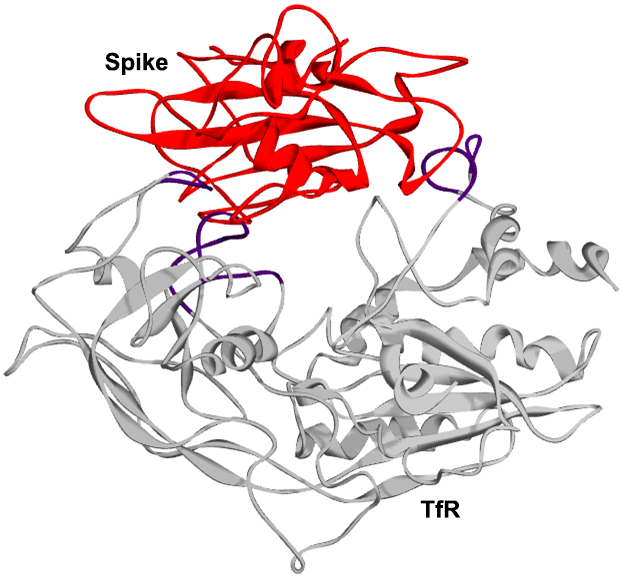

**Fig. S1. The 3D structure represents the complex of spike (red) and transferrin receptor (gray).** Known crystal structure of transferrin receptor (TfR) (PDB ID: 1CX8) and SARS-CoV-2-spike (PDB ID: 6LZG) were used for protein docking.

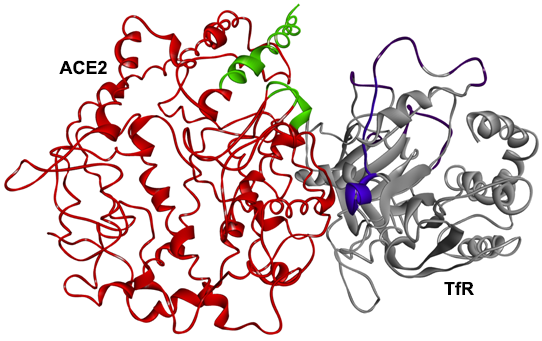

**Fig. S2. The 3D structure represents the complex of angiotensin-converting enzyme 2 (red) and transferrin receptor (gray).** Known crystal structure of transferrin receptor (TfR) (PDB ID: 1CX8) and angiotensin-converting enzyme 2 (ACE2) were used for protein docking.

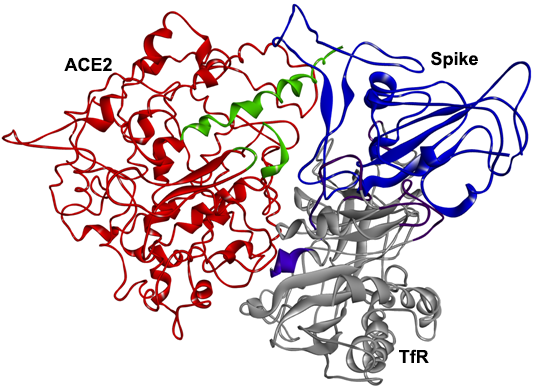

**Fig. S3. The 3D structure represents the transferrin receptor- angiotensin-converting enzyme 2-Spike complex.** Known crystal structure of transferrin receptor (TfR), angiotensin-converting enzyme 2 (ACE2), and spike were used for protein docking.

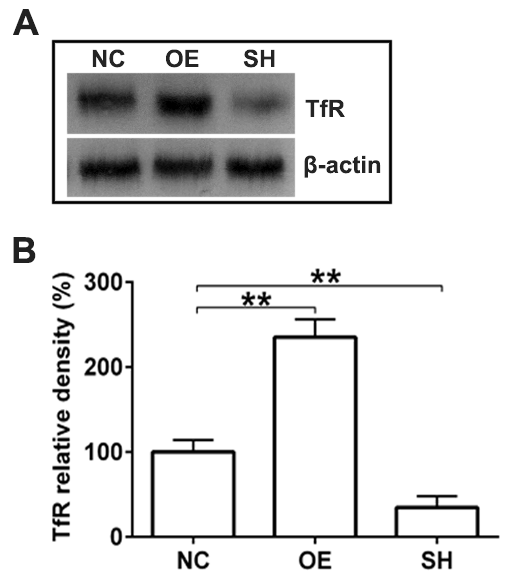

**Fig. S4.** **Construction of transferrin receptor overexpression and knockdown vectors.** **(A)** Transferrin receptor (TfR) levels in Vero E6 cells were determined by western blot (Lane 1: control (NC), Lane 2: overexpression (OE), Lane 3: knockdown (SH)). Quantification of western blot is shown **(B)**. Data represent mean ± SD of six independent experiments, ***p* < 0.01 by unpaired t-test.

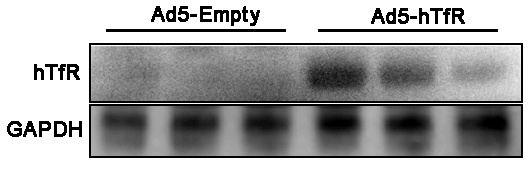

**Fig. S5. Validation of hTfR expression in lung tissue of mice.** TfR expression in lung tissue of mice transduced intranasally with empty AD5 (AD5-Empty) or Ad5-hTfR adenoviral vector was validated by western blot. GAPDH was used as loading control.

**Supplementary Table**

**Table S1. Designed peptides interfering with the interaction between transferrin receptor and spike.**

| Peptide | Sequence |
| --- | --- |
| SL8 | SKVEKLTL |
| QK8 | QDSNWASK |

**Table S2. Designed peptides interfering with the interaction between transferrin receptor and angiotensin-converting enzyme 2.**

| Peptide | Sequence |
| --- | --- |
| SL8 | SKVEKLTL |
| FG8 | FPFLAYSG |

**Supplementary references**

1	Tang, X. *et al.* Transferrin plays a central role in coagulation balance by interacting with clotting factors. *Cell Res* **30**, 119-132, doi:10.1038/s41422-019-0260-6 (2020).

2	Zhang, Z. *et al.* Mitochondrial DNA-LL-37 Complex Promotes Atherosclerosis by Escaping from Autophagic Recognition. *Immunity* **43**, 1137-1147, doi:10.1016/j.immuni.2015.10.018 (2015).

3	Wang, M. *et al.* Remdesivir and chloroquine effectively inhibit the recently emerged novel coronavirus (2019-nCoV) in vitro. *Cell Res* **30**, 269-271, doi:10.1038/s41422-020-0282-0 (2020).

4	De Meyer, S. *et al.* Lack of Antiviral Activity of Darunavir against SARS-CoV-2. *medRxiv*, 2020.2004.2003.20052548, doi:10.1101/2020.04.03.20052548 (2020).

5	Lu, S. *et al.* Comparison of SARS-CoV-2 infections among 3 species of non-human primates. *bioRxiv*, 2020.2004.2008.031807, doi:10.1101/2020.04.08.031807 (2020).

6	Sun, J. *et al.* Generation of a Broadly Useful Model for COVID-19 Pathogenesis, Vaccination, and Treatment. *Cell* **182**, 734-743 e735, doi:10.1016/j.cell.2020.06.010 (2020).
